# Action potential counting at giant mossy fiber terminals gates information transfer in the hippocampus

**DOI:** 10.1101/158444

**Authors:** Simon Chamberland, Yulia Timofeeva, Alesya Evstratova, Kirill Volynski, Katalin Tóth

**Affiliations:** Quebec Mental Health Institute, Department of Psychiatry and Neuroscience, Université Laval, 7 Quebec City, Quebec, Canada; Department of Computer Science, University of Warwick, Coventry, UK; Centre for Complexity Science, University of Warwick, Coventry, UK; University College London Institute of Neurology, University College London, London, UK

**Keywords:** information transfer, calcium, presynaptic, neurotransmitter release, information coding, action potential counting

## Abstract

Hippocampal mossy fibers have long been recognized as conditional detonators owing to prominent short-term facilitation, but the patterns of activity required to fire postsynaptic CA3 pyramidal neurons remain poorly understood. We show that mossy fibers count the number of spikes to transmit information to CA3 pyramidal cells through a distinctive interplay between presynaptic calcium dynamics, buffering and vesicle replenishment. This identifies a previously unexplored information coding mechanism in the brain.

## Main text

Neurons encode and transmit information in the frequency and temporal precision of action potentials (APs) they discharge ^1,2^. Presynaptic terminals are key elements involved in the translation of electrical signals to neurotransmitter release ^3^. During active states, several types of neurons fire in bursts. For example, hippocampal granule cells fire infrequently, but discharge bursts of APs with highly variable frequencies ^4,5^ However, how presynaptic mossy fiber bouton (MFB) terminals decode the frequency and the number of APs in the incoming bursts to transmit information remains poorly understood.

We first aimed to determine how AP transmission to CA3 pyramidal cells is encoded by the frequency and the number of APs discharged by granule cells. We recorded CA3 pyramidal cells in current-clamp and stimulated mossy fibers using trains of APs with the initial frequency of the first 5 stimuli delivered at 20 or 100 Hz and the last 3 stimuli fixed at 100 Hz. As expected AP firing by CA3 cells progressively increased during mossy fiber stimulation (Fig. 1a-c). The probability of observing the first postsynaptic spike sharply increased at the 6^th^ stimuli (Fig. 1d). Both the probability of CA3 pyramidal cell firing at the 6^th^ stimulus and the probability of observing the first AP were independent from the initial burst frequency (Fig. 1c,d). This suggests that AP transmission at MFB terminals is mainly determined by the number of spikes within the train and not by the average train frequency. Glutamate release from MFBs is greatly amplified during trains of stimuli ^6,7,8^, however how the frequency and number of stimuli are translated to specific patterns of glutamate release remains unknown. We varied the burst frequency and the number of stimuli to dissect the contribution of these two parameters. The 6^th^ evoked post synaptic current (EPSC) amplitude in a 5X20 Hz + 1X100 Hz burst was nearly identical to the 6^th^ EPSC amplitude of a pure 100 Hz train (Fig. 1e,f). Similarly, the 6^th^ EPSC amplitude of a 5X100 Hz + 1X20 Hz burst closely matched the amplitude of the 6^th^ EPSC in a 20 Hz train (Fig. 1g,h). This supports the idea that the average frequency of the train is not a determining factor of the rate of glutamate release. Instead, the number of preceding stimuli and the frequency of only the last stimulus appear to dictate the efficiency of synchronous glutamate release at the last 6^th^ spike. These data argue that MFB terminals use a counting logic. We confirmed that such counting logic was observed for any stimulus number between 2 – 10 (**Supplementary Fig. 1a,b**) and for frequencies between 10 and 100 Hz (**Supplementary Fig. 1c**).

**Figure 1.**
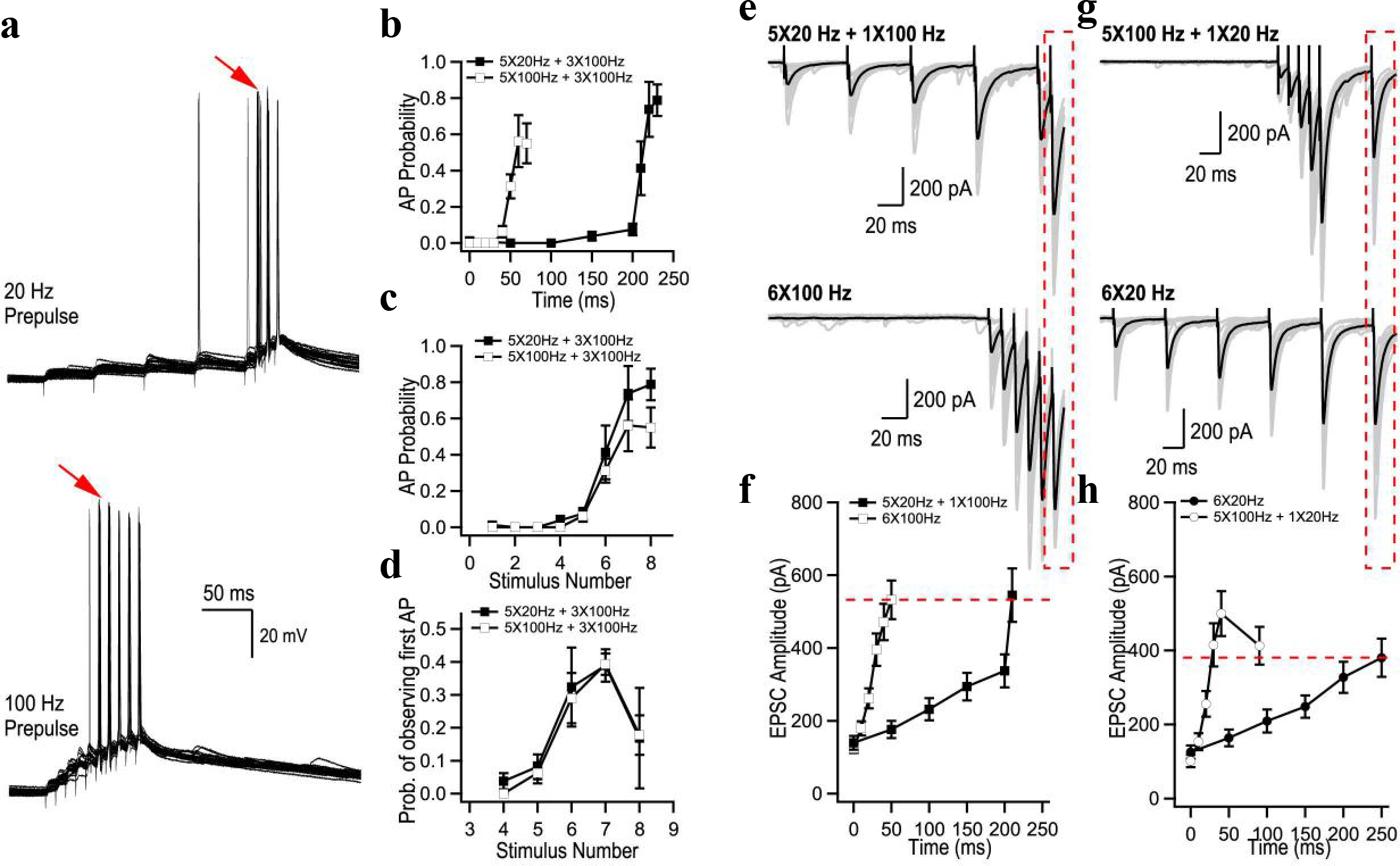
Transmission of information to CA3 pyramidal cells depends on the number of APs in bursts. (**a**) Representative current-clamp recordings from a CA3 pyramidal cell. The red arrows point to the 6^th^ stimulus in the trains where postsynaptic AP probability increased sharply. (**b, c**) CA3 pyramidal cell firing probability as a function of time (**b**) and of the number of APs (**c**) during two stimulation paradigms. (**d**) Probability of observing the first AP. Both AP probability and the probability of observing the first AP are mainly determined by the number of preceding APs but not by the average burst frequency (b-d, n = 5 cells). (**e-h**) Analysis of short-term facilitation at MFBs. (**e,g**) Representative trains of EPSCs recorded using indicated stimulation paradigms. Gray traces, single trials; black traces, the averages of 20 trials. (**f,h**) Summary graphs showing average EPSC amplitude as a function of time (n = 19 for 20 Hz; n = 15 for 5X20 Hz + 1X100 Hz; n = 25 for 100 Hz; n = 10 for 5X100 Hz+ 1X20 Hz).

To gather insights on the presynaptic determinants of the counting logic, we next performed fast whole-bouton two-photon random-access Ca^2+^ imaging using the low-affinity Ca^2+^ indicator Fluo-4FF to measure the dynamic modulation of presynaptic [*Ca*^2+^] during AP trains (Fig. 2). We found that the amplitude of AP-evoked Ca^2+^-fluorescence transients remained constant during AP bursts (Fig. 2b-e). This indicates that AP-evoked Ca^2+^ influx does not change during 20 Hz or 100 Hz stimulations and therefore, modulation of voltage-gated Ca^2+^ channel (VGCCs) activity is unlikely to contribute to short-term plasticity in MFB terminals. We next explored the presynaptic Ca^2+^ dynamics by direct fitting of the experimental traces using a nonstationary single compartment model^9,10^ (Fig. 2b,c, Online Methods). The model, which incorporated three major endogenous Ca^2+^ buffers known to be present in MFBs (calbindin-D_28K_ (CB), calmodulin (CaM), and ATP) provided close fits of the experimental data (Fig. 2b,c and **Supplementary Fig. 2**). It is noteworthy that a similar model with a single fast high affinity endogenous buffer^11^ could not replicate the Ca^2+^ imaging data (**Supplementary Fig. 3**). The fitting allowed us to estimate Ca^2+^ removal rate in our experimental conditions (*k_rem_* range 0.2 - 0.7 ms^-1^), which was in close agreement with previous estimates obtained with high affinity Ca^2+^ indicator Fluo-4^9^.

**Figure 2.**
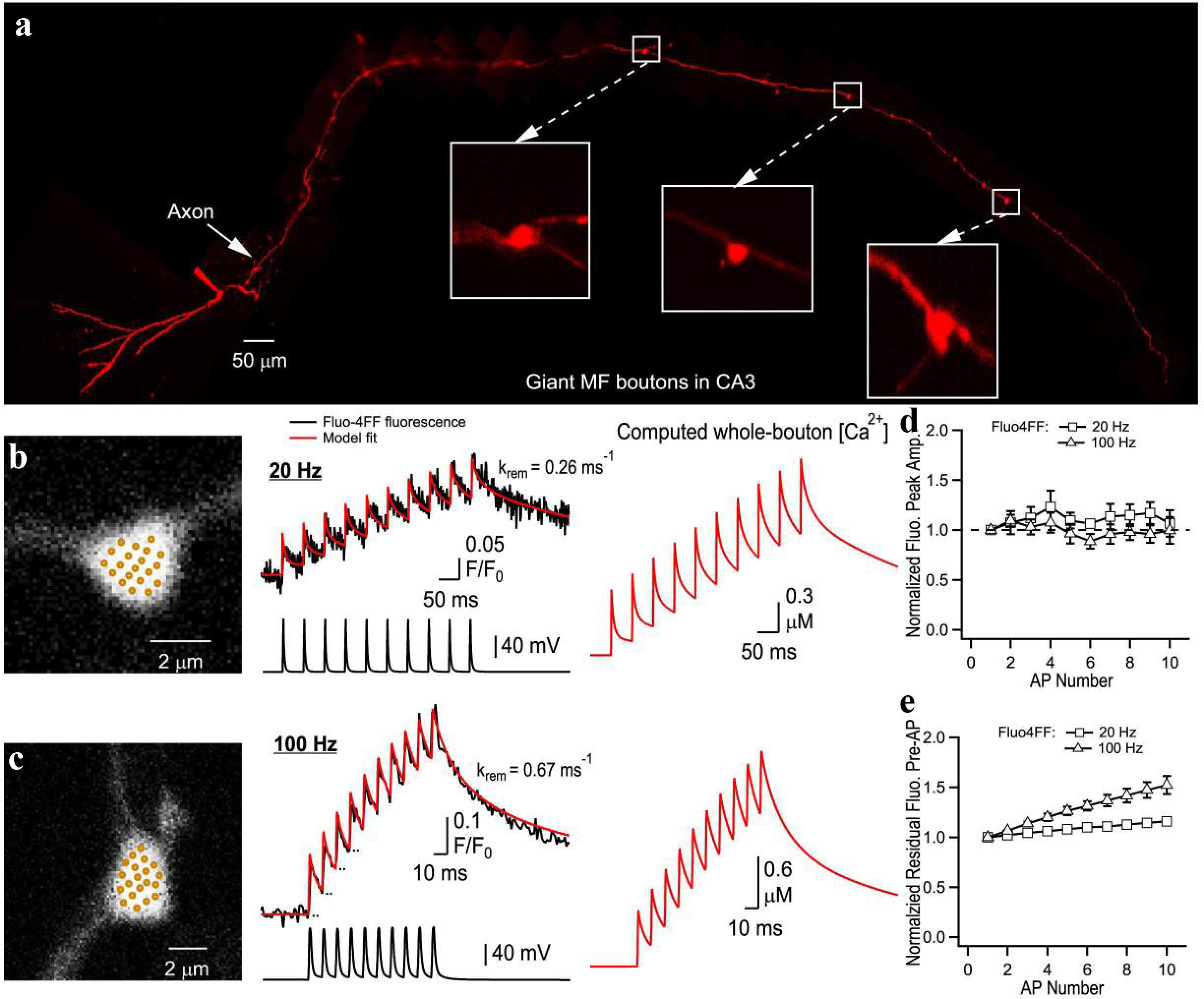
Ca^2+^ dynamics in MFB terminals. (**a**) Montage of multiple two-photon Z-stack maximal projections showing the axon anatomy of a typical granule cell filled with a morphological tracer AlexaFluor 594. The intact axon was followed to the CA3 region, where it formed giant MFB terminals (insets). (**b,c**) Representative measurements of Ca^2+^ dynamics in two MFBs. Left, bouton morphology, dots indicate recording positions for random-access two-photon Ca^2+^ imaging. Middle, corresponding whole bouton Ca^2+^ Fluo-4FF fluorescence elevations in response to 20 Hz and 100 Hz AP stimulations (black traces, average of 140 and 133 sweeps, respectively). The red curves represent the non-stationary single compartment model fit corresponding to Δ[Ca^2+^]_total_ = 33.3 μM, model-predicted values of *k_rem_* are shown for each bouton. Right, corresponding AP-evoked whole-bouton [Ca^2+^] transients computed using the non-stationary model. (d) Normalized peak amplitude of AP-evoked Fluo-4FF fluorescence as a function of AP number (n = 7 MFBs for both 20 Hz and 100 Hz stimulation).

To understand whether the interplay between presynaptic Ca^2+^ dynamics and endogenous Ca^2+^ buffering can lead to AP counting, we performed quantitative modelling of AP-evoked Ca^2+^ influx, buffering and diffusion, and glutamate release in MFBs. The three-dimensional model incorporated key ultrastructural and functional properties of MFBs including multiple release sites, experimentally constrained presynaptic Ca^2+^ dynamics and loose coupling between VGCCs and vesicular release sensors ^9,11-13^ (Fig. 3a and Online Methods). The simulation unit, which represented a part of MFB with a single release site, was modelled as a parallelepiped of size 0.5 μpm × 0.5 μm × 0.79 μm with a single VGCC cluster in the middle of the bottom base (Fig. 3a). As in the case of the single compartment model we assumed the presence of three major MFB endogenous Ca^2+^ buffers: CB, ATP and CaM. At physiological conditions CaM is known to be distributed between membrane-bound and mobile states, and this distribution is regulated by intracellular [Ca^2+^]^14,15,16^. We first considered a limiting case of ‘Mobile CaM’ model. We simulated spatial MFB Ca^2+^ dynamics in response to bursts of APs and used the obtained [*Ca*^2+^] transients at the release site (90 nm away from the VGCC cluster, (Fig. 3b) to perform simulations of vesicular release using a Monte Carlo implementation of Ca^2+^-activated vesicle fusion model^12^ (Fig. 3c and **Supplementary Fig. 4**). To account for vesicle replenishment during AP bursts we included a vesicle replenishment step in the model and experimentally constrained the replenishment rate constant (*k_rep_* = 20 s^-1^) (**Supplementary Fig. 5**). We found that ‘Mobile CaM’ model indeed replicated the AP counting during mixed 20 Hz and 100 Hz AP trains (Fig. 3d,e and **Supplementary Fig. 6**). What mechanisms underlie the counting logic? The model predicted that the peak values of Ca^2+^ transients ([*Ca*^2+^]_*peak*_) were gradually augmented during

**Figure 3.**
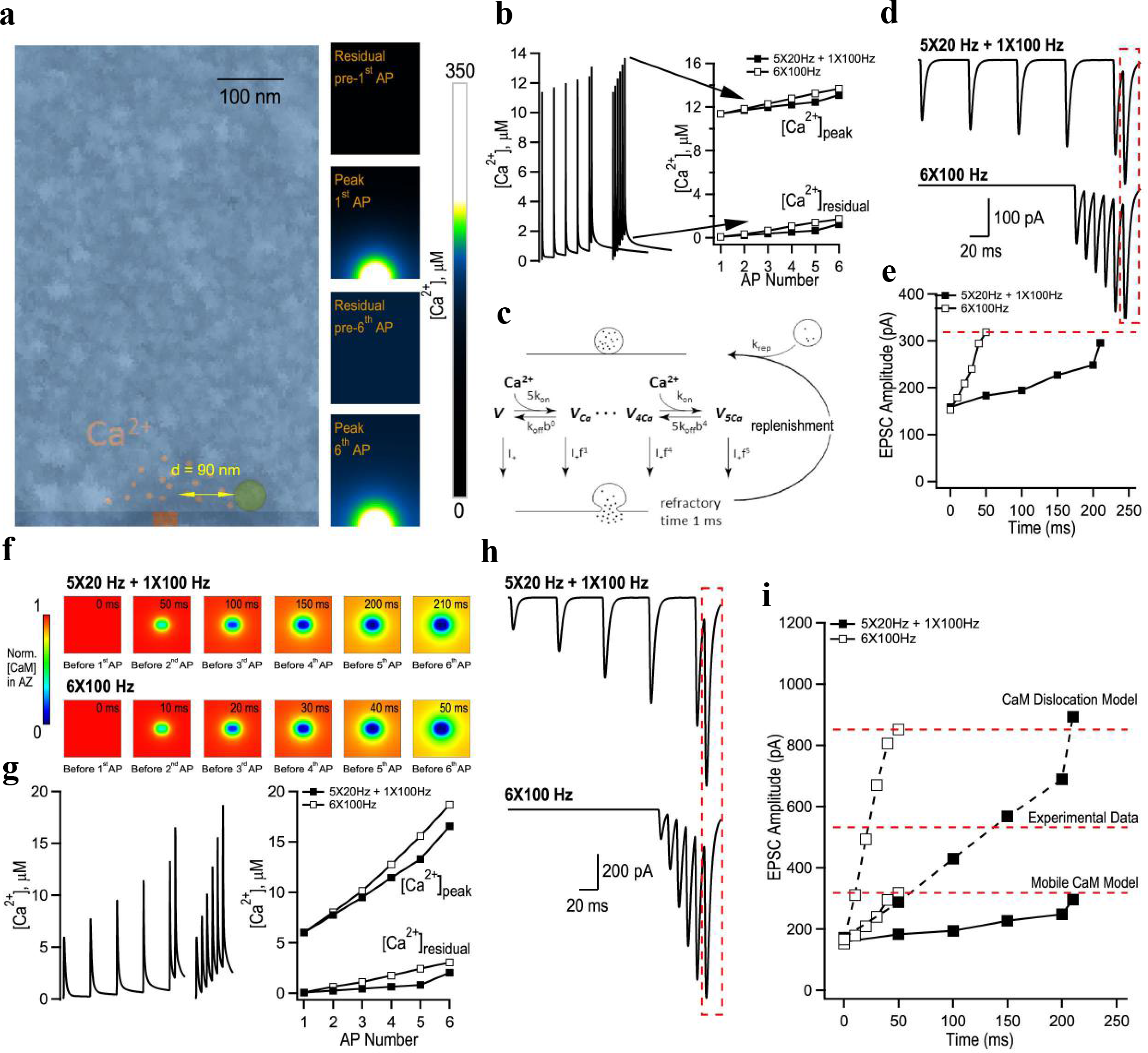
Modelling of evoked presynaptic Ca^2+^ dynamics and vesicular release reveals a plausible mechanism for AP counting logic in MFB terminals. (**a**) Left, geometry of a parallelepiped-shaped modelling unit (0.5 μm × 0.5 μm × 0.79 μm) used in VCell simulations representing part of a MFB containing a single active zone (AZ) with a 40 nm × 80 nm VGCC cluster, overlaid with a representative electron microscopy image that depicts part of a MFB with a single AZ (see Online methods for details). Right, snapshots of VCell-computed spatial [Ca^2+^] profiles in the central XZ modelling unit plane during 6X100 Hz AP stimulation in the limiting case of ‘Mobile CaM’ model. (**b**) Left, VCell-computed [Ca^2+^] transients at the release site located at *d* = 90 nm away from the VGCC cluster during 6X100 Hz and 5X20 + 1X100 AP trains. Right, plot of corresponding residual ([Ca^2+^]_residual_) and peak ([Ca^2+^]_peak_) before and after each AP in the trains (**c**) Schematics of the modified allosteric model of Ca^2+^-driven vesicle release and replenishment ^12^. (**d**) Simulated EPSCs, average of M = 60,000 Monte Carlo runs for each paradigm scaled for RRP of size m = 125. (**e**) Summary graph showing simulated EPSC amplitude as a function of time for the two stimulation paradigms shown in (d). (**f**) Snapshots of spatial distribution of normalized total [CaM] (which accounts for all CaM molecules irrespective of their Ca^2+^ binding state) in the AZ plane, illustrating progressive dislocation of CaM from the membrane during AP stimulation predicted by ‘CaM dislocation model’ ^14^ (see Online methods for details). (**g**) VCell-computed [Ca^2+^] transients at the release site during 6X100 Hz and 5X20 Hz + 1X100 Hz AP trains and (**h**) corresponding simulated EPSCs for the case of ‘CaM dislocation’ model. (**i**) Summary graph showing that experimentally observed short-term facilitation levels are likely to be explained by joint contribution of the two limiting cases represented by ‘Mobile CaM’ (low facilitation) and by ‘CaM dislocation’ (high facilitation) models that both allow AP counting logic.

AP bursts which was mainly attributed to the increase in residual [Ca^2+^]_*residual*_, and was mostly independent of the stimulation frequency (Fig. 3b). This argues that EPSC facilitation predicted by ‘Mobile CaM’ model was due to [*Ca*^2+^]_*residual*_, accumulation and not due to endogenous Ca^2+^ buffer saturation (which normally leads to a progressive increase of the amplitudes of individual AP-evoked Ca^2+^ transients [*Ca*^2+^]_*amp*_ = [*Ca*^2+^]_*peak*_ – [*Ca*^2+^]_*residual*_ ^13^). Indeed, the model revealed that fast and low affinity CaM N-lobe did not show progressive saturation. However, slower and high affinity buffers CB and CaM C-lobe did saturate during AP bursts (**Supplementary Fig. 7**). This at first sight contradictory observation was fully in line with the dominant effect of CaM N-lobe on release site Ca^2+^ dynamics and vesicle fusion^14^ (**Supplementary Fig. 8**).

Although ‘Mobile CaM’ model replicated AP counting the overall level of EPSC facilitation predicted by this model was ∼ 40% lower than the experimentally observed values (Fig. 1f, Fig. 1h). Therefore, we considered another limiting case, ‘CaM dislocation’ model. In this model (described in detail in our previous publication ^14^) we considered that CaM was initially bound to the presynaptic membrane via interaction of its C-lobe with neuromodulin and with other IQ-motif presynaptic membrane proteins (e.g. VGCCs)^14,15,16^. The model assumed that Ca^2+^ binding by the CaM C-lobe during AP bursts led to dissociation of CaM from its membrane binding partners and thus resulted in a stimulation-dependent reduction of Ca^2+^ buffering capacity in the AZ (Fig. 3f). This in turn led to progressive increase of [Ca^2+^] transients at the AZ (Fig. 3g) and to facilitation of EPSCs (Fig. 3h). The progressive reduction of AZ Ca^2+^ buffering capacity predicted by the model did not depend on the frequency of AP bursts. Thus, ‘CaM dislocation’ model also supported the counting logic at MFB terminals. In contrast to ‘Mobile CaM’ model the dislocation model predicted substantial increase of AP-evoked [*Ca*^2+^]_*amp*_ at the release site which resulted in stronger EPSC facilitation (Fig. 3g,i). Overall, the experimentally observed level of EPSC facilitation in MFB terminals is likely to be attributed to a joint contribution of the ‘Mobile CaM’ and ‘CaM dislocation’ limiting cases (Fig. 3i). Interestingly, the effect of somewhat stronger augmentation of [*Ca*^2+^]_*peak*_ on vesicular release at higher frequencies was compensated in both models by the lower vesicle occupancy at the release site during high frequency stimulation (**Supplementary Fig. 10**). This indicates that frequency-dependent differences in release site occupancy also contribute to the counting logic behavior of MFBs.

We aimed to understand how granule cells generate CA3 pyramidal cells firing, the first relay of information in the hippocampus ^17^ This question is important because discharge of a single AP by a single CA3 pyramidal cell has dramatic network consequences such as initiation of sharp-wave ripples ^18^. We demonstrate that MFB count the number of spikes to transmit information to CA3 pyramidal cells independently of the average spike frequency. Our results argue that MFB counting logic can be explained by (i) accumulation of [*Ca*^2+^]_*residual*_, which is largely independent of stimulation frequency due to relatively slow Ca^2+^ removal rate *k_rem_*; (ii) loose coupling between VGCCs and vesicular release sensors which leads to moderate changes of [*Ca*^2+^]_*peak*_ at release site (in the range of 10 μM - 15μM) that is efficiently modulated by changes in [*Ca*^2+^]_*residual*_ (∼ 2 μM); (iii) the dominant role of CaM N-lobe on fast Ca^2+^ buffering at the release sites; (iv) possible contribution of stimulation-dependent reduction of Ca^2+^ buffering capacity in the AZ due to CaM dislocation, and; (v) more efficient vesicle replenishment at lower frequencies. All of the above elements are uniquely combined in MFBs to allow AP counting, an information coding strategy which contrasts rate and temporal coding described in other types of synapses ^2^.

## Online Methods

### Hippocampal slice preparation

Experiments involving the use of animals were performed in accordance with guidelines provided by the Animal Protection Committee of Laval University. Acute hippocampal slices from P17 – P25 male rats were prepared according to accepted procedures^7^. First, the animals were anesthetized with isoflurane. The brain was extracted and immersed in an oxygenated cutting ACSF solution maintained at 4 °C. The cutting ACSF solution contained (in mM): NaCl 87, NaHCO_3_ 25, KCl 2.5, NaH_2_PO_4_ 1.25, MgCl_2_ 7, CaCl_2_ 0.5, glucose 25 and sucrose 75 (pH = 7.4, 330 mOsm). The brain was then dissected according to instructions for optimal preservation of the hippocampal mossy fibers^19^. The brain hemispheres were glued on the specimen disk of a Leica VT1000S vibratome and submerged in cutting ACSF solution. Slices (300 μM) were cut and transferred to an oxygenated and heated (32 °C) ACSF solution containing (in mM): NaCl 124, NaHCO_3_ 25, KCl 2.5, MgCl_2_ 2.5, CaCl_2_ 1.2 and glucose 10 (pH = 7.4, 300 mOsm). Slices were left to recover for 30 minutes at 32 °C. Slices were then left at room temperature. Experiments were started one hour after the slicing procedure.

### Whole-cell patch-clamp recording

Hippocampal slices were maintained under a nylon mesh in a recording chamber under an upright microscope. The slice was perfused with oxygenated warmed recording ACSF solution, containing (in mM): NaCl 124, NaHCO_3_ 25, KCl 2.5, MgCl_2_ 2.5, CaCl_2_ 1.2 and glucose 10. The solution was oxygenated by bubbling a gas mixture composed of 95% O2 and 5% CO2. Temperature was maintained at 32 ± 1 °C throughout all experiments. The perfusion rate was adjusted to a constant 2 ml/min. Visually-guided whole-cell patch-clamp recordings were obtained from CA3 pyramidal cells with a solution containing: K-gluconate 120, KCl 20, HEPES 10, MgCl_2_ 2, Mg_2_ATP 2, NaGTP 0.3, phosphocreatine 7, EGTA 0.6 (pH = 7.2, 295 mOsm). Borosillicate glass electrodes had a resistance of 3 – 5 MΩ for CA3 pyramidal cell recordings. After obtaining a stable whole-cell configuration, CA3 pyramidal cells were held in voltage-clamp or in current-clamp. Voltage-clamp recordings were performed at −70 mV. Current-clamp recordings were performed at the resting membrane potential of the CA3 pyramidal cells (-70 ± 5 mV). Minimal stimulation of mossy fibers was performed using an electrode positioned in the stratum lucidum and connected to a constant current stimulus isolator (A360, WPI, Florida, USA). The pipette was gently moved in the stratum lucidum until large, fast and facilitating EPSCs could be recorded. The stimulation intensity was then decreased to achieve conditions in which both failures and successes could be observed. To confirm the mossy fiber identity of the recorded EPSCs or EPSPs, DCG-IV (1 μM) was applied in the end of a subset of experiments. Recordings in which the postsynaptic response was decreased by at least 80% were conserved for further analysis. Electrophysiological data was acquired with Molecular Devices equipment (Axopatch 200B amplifier and Digidata 1322A, or MultiClamp 700B amplifier with Digidata 1440A) and the Clampex suite. The electrophysiological data was low-pass filtered at 2 kHz, digitized at 10 kHz and recorded on a personal computer. For calcium imaging experiments, whole-cell patch-clamp recordings were obtained from granule cells with the solution described above, but lacking EGTA. This patch solution was supplemented with 40 μM of the morphological dye Alexa-594 and 375 μM of the low-affinity calcium indicator Fluo-4FF. Granule cells were held in the current-clamp mode at their resting membrane potential. Action potentials were evoked by brief current injections (2 ms, 1 – 1.5 nA) in trains of 10 APs, at either 20 Hz or 100 Hz. Glass electrodes used for whole-cell recordings from granule cells had a resistance between 4 – 7 MΩ.

### Random-access two-photon calcium imaging

Following diffusion for at least 1 hour of the fluorophores in the granule cell, the axon was tracked to the CA3 region^7,9^. Giant MF boutons were unequivocally identified in the CA3 region based on their morphology imaged with the AlexaFluor-594 fluorescence. 20 sites evenly dispersed on the whole bouton were recorded quasi-simultaneously, yielding an imaging speed of 950 Hz. This recording paradigm allowed a good compromise between signal to noise ratio of the signal and the temporal resolutions, and therefore enabled recording calcium elevations generated by high-frequency firing of APs. The very low-affinity Ca^2+^ indicator Fluo-4FF proved critical to resolve high-frequency bursts of APs evoked at 100 Hz without indicator saturation. We used a custom built random-access two-photon microscope^7,20^. A two-photon titanium:sapphire laser (Chameleon Ultra II, Coherent) tuned at 800 nm provided the light source (80 MHz, 140 fs pulse width and with an average power >4 W). The laser beam was redirect by a pair of acousto-optic deflectors (AODs; A-A Opto Electronics) to enable random access over the field of view. The laser beam was focused on the brain slice through a high NA water-immersion objective (25X objective, with a NA = 0.95). Transmitted photons passed through a high-numerical aperture oil condenser (NA = 1.4) and were low-pass filtered at 720 nm. Photons were separated by a dichroic mirror (580 nm) to independently collect red and green photons. Photons were then band-pass filtered at 500-560 nm for the green channel and 595-665 nm for the red channel. Both the red and the green photons were collected simultaneously. Collection of photons was performed using a pair of AsGaP photomultiplier tubes (H7422P-40, Hamamatsu) located close to the recording chamber. The laser and the acquisition system were controlled by a Labview custom-made software^20^.

### Analysis of electrophysiological and calcium imaging data

Electrophysiological data was analyzed in Clampfit and in Igor Pro. AP probability was calculated from 20 sweeps. To avoid inducing long-term plasticity, sweeps were evoked every 30 seconds. EPSC amplitude was measured from the average trace obtained from 20 sweeps. Calcium elevations recorded in giant MF terminals were exported to Excel database. The ΔG/G ratio was calculated for all trials and trials (50 – 140) were averaged together. The peak Ca^2+^amplitude for individual calcium transients was determined from baseline to peak. In all figures, symbols show the mean and the error bars indicate the SEM.

### Non-stationary single compartment model of presynaptic Ca^2+^ dynamics

Experimental Ca^2+^ fluorescence traces were analysed using a non-stationary single compartment model^9,10^, which assumes spatial homogeneity of [Ca^2+^] in the nerve terminal. The model is described by the following system of differential equations:

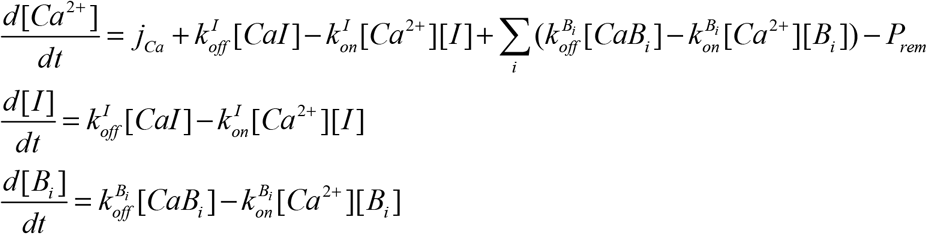

where the square brackets denote concentrations, and the superscript indices of the reaction rate constants denote endogenous Ca^2+^ buffers *B_i_* or the Fluo-4FF indicator *I*. The action potential-dependent Ca^2+^ influx time course *j_Ca_* was approximated by the Gaussian function 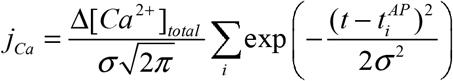 where 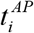 denotes the times of peaks of Ca^2+^ currents during each action potential. The use of the low affinity Ca^2+^ indicator Fluo-4FF (*K_d_ =* 9.7 μM) did not allow us to estimate resting [Ca^2+^]_*rest*_ reliably, which in turn prevented the numerical estimation of the total volume averaged presynaptic Ca^2+^ entry Δ[Ca^2+^]_*total*_. Therefore we used previous estimates for both [Ca^2+^]_*rest*_ = 75 nM and Δ[Ca^2+^]_*total*_ =50 μM obtained with high affinity Ca^2+^ indicators ^9,21^ Because in our experimental conditions [Ca^2+^]_*ext*_ = 1.2 mM (in comparison to [Ca^2+^]_*ext*_ = 2 mM in ref. ^9^) we reduced Δ[Ca^2+^]_*total*_ determined in ref.^9^ by a factor of 1.5 based on the dependency of VGCC conductance on [Ca^2+^]_*ext*_. Ca^2+^ removal was approximated by a first-order reaction *P_rem_* = *k_rem_*([Ca^2+^]-[Ca^2+^]_*rest*_). We assumed that a MFB terminal contains three endogenous buffers ATP, CB and CaM. The complete set of model parameters and Ca^2+^ binding reactions is specified in **Supplementary Table 1**. The model was numerically solved using the adaptive step-size Runge-Kutta algorithm. The model operated with only two adjustable (free) parameters: the unknown ratio between resting Fluo-4FF fluorescence signal and the background fluorescence and Ca^2+^ removal rate *k_rem_*. Both parameters were constrained by a straightforward fitting procedure that would match the calculated and experimental fluorescence profiles.

### Spatial VCell model of MFB Ca^2+^ dynamics

Three-dimensional modeling of AP-evoked presynaptic Ca^2+^ influx, buffering, and diffusion was performed in the Virtual Cell (VCell) simulation environment (http://vcell.org) using the fully implicit adaptive time step finite volume method on a 10 nm meshed geometry. A simulation unit, representing part of a MFB terminal with a single active zone (AZ), was modeled as a parallelepiped of size *x* = 0.5 μm, *y* = 0.5 *μ*m and *z* = 0.79 *μ*m. The AZ was located in the XY base (*z* = 9.79 *μ*m) and contained a single rectangular VGCC cluster of dimensions 40 nm × 80 nm placed in the center of the AZ. The size of XY base corresponded to the average distance among different AZs in MFB terminals (0.5 *μ*m)^22^. The height of the simulation unit was adjusted to *z* = 0.79 *μ*m in order to match the magnitude of local VGCC-mediated Ca^2+^ influx at the AZ (see below) to the value of experimentally estimated Δ[*Ca*^2+^]_*total*_ = 33.3 μM. We assumed that 28 VGCCs were evenly distributed within the VGCC cluster ^11,13^. The average AP-evoked Ca^2+^ current was simulated using the five-state VGCC gating kinetic model in MFB using the NEURON simulation environment^14,23^ and the experimentally determined MFB AP waveform^11^, which was considered to be constant during burst of APs. Ca^2+^ extrusion by the bouton surface pumps (excluding the AZ) was approximated by a first-order reaction: *j_extr_* = *k_extr_*([*Ca*^2+^]-[*Ca*^2+^]_*rest*_) ^10,24^ located at the XY parallelepiped base opposite to the AZ; *k_extr_* was calculated using the experimentally constrained single-compartment model average Ca^2+^ removal rate (*k_rem_* = 400 s^-1^) as 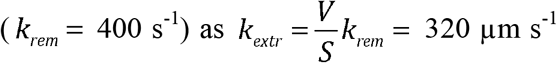 (where *V* is the volume of the simulation unit and *S* is the area of the XY base). In the case of ‘Mobile CaM’ model we assumed [*CaM*]_*total*_ = 150 μM as was estimated in **Supplementary Fig. 8a**. In the case of ‘CaM dislocation’ model we assumed that all CaM molecules were located within a single 10 nm layer of VCell voxels adjacent to the AZ plasma membrane (i.e. at the 0.5 μm × 0.5 μm bottom base of the simulation unit). Concentration of CaM was 3 molecules / 10 nm × 10 nm × 10 nm voxel, as estimated in **Supplementary Fig. 8c**. The details of ‘CaM dislocation’ model are described in our previous publication (ref. 29). Briefly, we assumed that upon binding of two Ca^2+^ ions by the C-lobe a CaM molecule can irreversibly dissociate from the plasma membrane (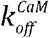 = 650 s^-1^, ref. 29) and freely diffuse in the cytosol.

### Modeling of Ca^2+^-triggered synaptic vesicle fusion

We assumed that the vesicular Ca^2+^ release sensor was located at coupling distance *d* = 90 nm from the edge of VGCC cluster (Fig. 3a). To simulate glutamate release we used [*Ca*^2+^](*t*) profiles obtained in VCell at this location for each specific AP firing pattern in Monte Carlo simulations (implemented in MATLAB) based on the six-state allosteric Ca^2+^ sensor model^12^ (Fig. 3c). The model also contained a stochastic re-priming step, which was preceded by a short refractory period (1 ms) immediately after vesicle fusion. The model parameters were: *k_on_* = 100 μM^-1^ s^-1^, *k_off_* = 4 × 10^3^ s^-1^, *b* = 0.5, *f* = 31.3, *I*_+_ = 2 × 10^-4^ s^-1^. The re-priming rate *k_rep_* = 20 s^-1^ was constrained using the experimental data for 50 stimuli applied at 100 Hz (**Supplementary Fig. 5**). For each stimulation paradigm, we performed 60,000 independent Monte Carlo runs with a time step *dt* = 10^-6^ s and thus determined distribution for the vesicle fusion time during AP burst. Simulated EPSC response was calculated as 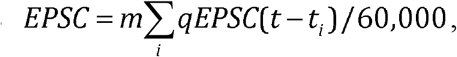, where *m*= 125 is the average RRP size and *qEPSC(t)* is the average quantal EPSC which was determined using voltage-clamp recoding during the asynchronous phase of release (200 – 300 ms after the last AP in the burst).

## Acknowledgements

The study was supported by CIHR (MOP-81142) and NSERC (RGPIN-2015-06266) grants (K.T.), NSERC and CTRN PhD fellowships (S.C.) and by the Medical Research Council and the Wellcome Trust (K.V.). We are grateful to D.M. Kullmann for reading the manuscript and providing feedback.

